# Complement factor C1q mediates chronic neuron loss and inflammation post-brain injury

**DOI:** 10.1101/2020.05.29.120220

**Authors:** Stephanie S Holden, Oumaima Aboubakr, Bryan Higashikubo, Frances S Cho, Andrew H Chang, Allison Morningstar, Vidhu Mathur, Logan J Kuhn, Poojan Suri, Sethu Sankaranarayanan, Yaisa Andrews-Zwilling, Eleonora Aronica, Ted Yednock, Jeanne T Paz

## Abstract

While traumatic brain injury (TBI) acutely disrupts the cortex, most TBI-related disabilities reflect secondary injuries that accrue over time. The thalamus is a likely site of secondary damage because of its reciprocal connections with the cortex. Using a mouse model of cortical injury that does not directly damage subcortical structures, we found a chronic increase in C1q expression specifically in the corticothalamic circuit. Increased C1q expression co-localized with neuron loss and chronic inflammation, and correlated with altered cortical rhythms. Blocking C1q counteracted most of these outcomes, suggesting that C1q is a disease modifier in TBI. Since the corticothalamic circuit is important for sensory processing, attention, cognition, and sleep, all of which can be impaired by TBI, this circuit could be a new target for treating TBI-related disabilities.

## INTRODUCTION

Traumatic brain injury (TBI) affects about 69 million people worldwide every year (*1*) and can lead to cognitive dysfunction, difficulty with sensory processing, sleep disruption, and the development of epilepsy. Most of these adverse health outcomes develop months or years after TBI and are caused by indirect secondary injuries that result in long-term consequences of the initial impact (*2*). Because the primary injury is essentially irreversible, understanding where, when, and how secondary injuries develop is crucial for preventing or treating disability following TBI.

The cortex is often the site of primary injury because it sits directly beneath the skull, and is an integrated part of many larger circuits, including the cortico-thalamo-cortical loop. This circuit is important for sensory processing, attention, cognition, and sleep, all of which can be impaired by TBI (*3*). The thalamus itself, though not acutely injured in TBI, experiences secondary injury, presumably because of its long-range reciprocal connections with the cerebral cortex (*4–8*). Structural changes in the thalamus have been implicated in a number of long-term TBI-related health outcomes, including fatigue and cognitive dysfunction (*9–10*), and patients with TBI display secondary and chronic neurodegeneration and inflammation in thalamic nuclei (*11–12*).

Chronic neuroinflammation is a common feature of secondary injury sites (*13*). But most attempts to improve post-TBI cognitive outcomes with broad anti-inflammatory agents have failed (*13–14*), likely because there are many inflammatory pathways that play both protective and pathogenic roles at different times (*15*). A potential mediator of post-TBI inflammation and injury is the complement pathway, which is activated in the peri-injury area of brain lesions in both humans and rodents (*16–18*). Complement activation contributes to inflammation and neurotoxicity in central nervous system injury and is increased in human brains afflicted with injury, epilepsy, and Alzheimer’s disease (*18–22*). Aberrant activation of C1q, the initiating molecule of the classical complement cascade, can trigger elimination of functioning synapses and contribute to the progression of neurodegenerative disease (*23*). On the other hand, C1q is involved in normal synapse pruning during development (*24*) and the complement system plays an important part in brain homeostasis by clearing cellular debris and protecting the central nervous system from infection (*19*).

Here, we investigated the role of classical complement protein C1q in post-TBI impairment of the corticothalamic circuit, with a particular emphasis on the timing and location of C1q expression. We used a mouse model of mild TBI, and monitored neurophysiological changes in the corticothalamic circuit via cellular electrophysiology and wireless cortical recordings in freely behaving mice up to four months post-TBI.

## RESULTS

### Secondary C1q expression coincides with chronic inflammation, neurodegeneration, and synaptic dysfunction in the thalamus

To determine the secondary, long-term effects of TBI, we induced a mild cortical impact injury to the right primary somatosensory cortex (S1) of adult mice (Figure 1A), and assessed the impact on their brains three weeks later. This period corresponds to the latent phase in humans, when the brain is undergoing adaptive and maladaptive changes (*25*). We determined neuron count and gliotic inflammation in the corticothalamic circuit by immunofluorescent staining of coronal brain sections with markers of neurons (NeuN) and of glial inflammation (C1q, complement pathway; GFAP, astrocytes; Iba1, microglia/macrophages) (Figure 1C-E). Three weeks post-surgery, TBI mice had significantly higher GFAP, C1q, and Iba1 expression in the peri-TBI S1 cortex, the functionally connected ventrobasal thalamus (VB), and the reticular thalamic nucleus (nRT) than sham mice (Figure 1B-E). Inflammation occurred within 24 hours after injury in the cortex, while the functionally connected nRT and VB displayed glial changes around five days later (not shown), suggesting secondary thalamic inflammation. We also saw increased expression of similar inflammatory markers in thalamic tissue from human TBI patients, confirming that thalamic inflammation is a consequence of TBI in humans too (Figure S1). We conclude that a chronic inflammatory process, secondary to the injury and characterized by C1q expression, occurs in the thalamus.

**Figure 1.**
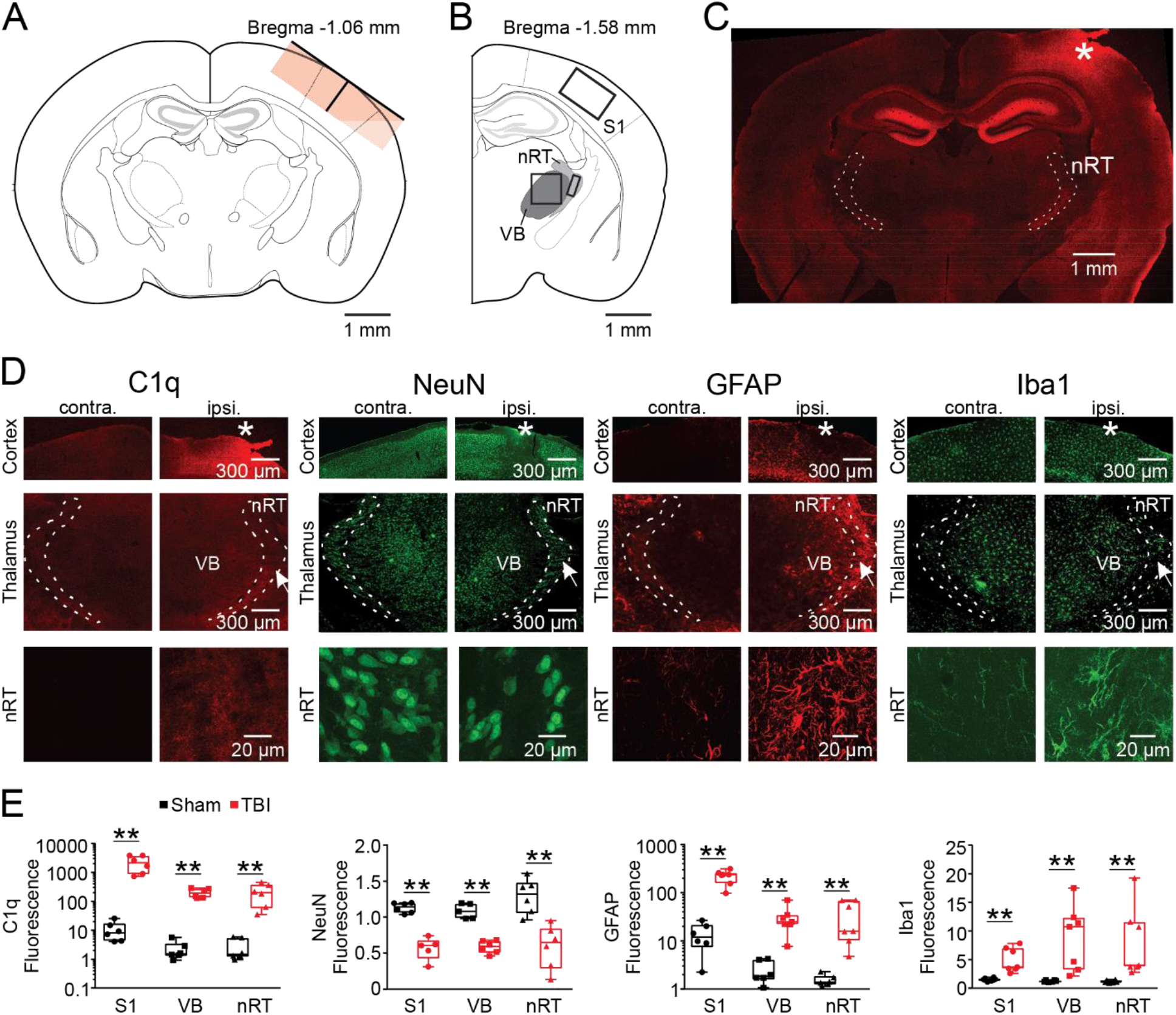
The injured cortex and functionally connected thalamus show chronic inflammation and neuron loss three weeks after TBI. A, B) Schematic of a mouse brain coronal section showing the site and depth of the controlled cortical impact (A) and the location of the S1 cortex and nRT and VB thalamic regions (B). The impactor has a diameter of 3 mm and the impact was delivered at a depth of 0.8 mm to the right somatosensory cortex. C) Representative coronal brain section from a TBI mouse stained for C1q. C1q expression in the hippocampus is typical of physiological conditions. D) Close-up images of S1 (top), VB and nRT (middle), and confocal images of nRT (bottom) stained for C1q, neuronal marker NeuN, GFAP, a marker for astrocytes, and Iba1, a marker for microglia/macrophages. Injury site in the right S1 cortex is marked by an asterisk. Arrow in nRT indicates location of confocal image. Scale bars, 300 μm (top/middle) and 20 μm (bottom). E) Quantification of fluorescence ratios between ipsilateral and contralateral regions in sham and TBI mice. Data represent all points from min to max, with a Mann-Whitney test and α = 0.05 (*p < 0.05, **p < 0.01). Analysis includes between five and seven mice per group (n = three sections per mouse, one image per region).

Glial inflammation was associated with significant neuronal loss in the thalamic region, particularly in the nRT (Figure 1D-E, Figure 2A), which receives the majority of its glutamatergic inputs from the cortex (*26–27*). The nRT of TBI mice had significantly fewer neurons than sham mice, particularly in the region of the nRT that receives most of its excitatory inputs from the injured somatosensory cortex (*26–29*) (Figure 2B-C). This result suggests that the inflammation, which may be initiated by retrograde axonal degeneration, follows the long-range, corticothalamic circuit, and marks its two ends: the injured cortex and the connected thalamus.

**Figure 2.**
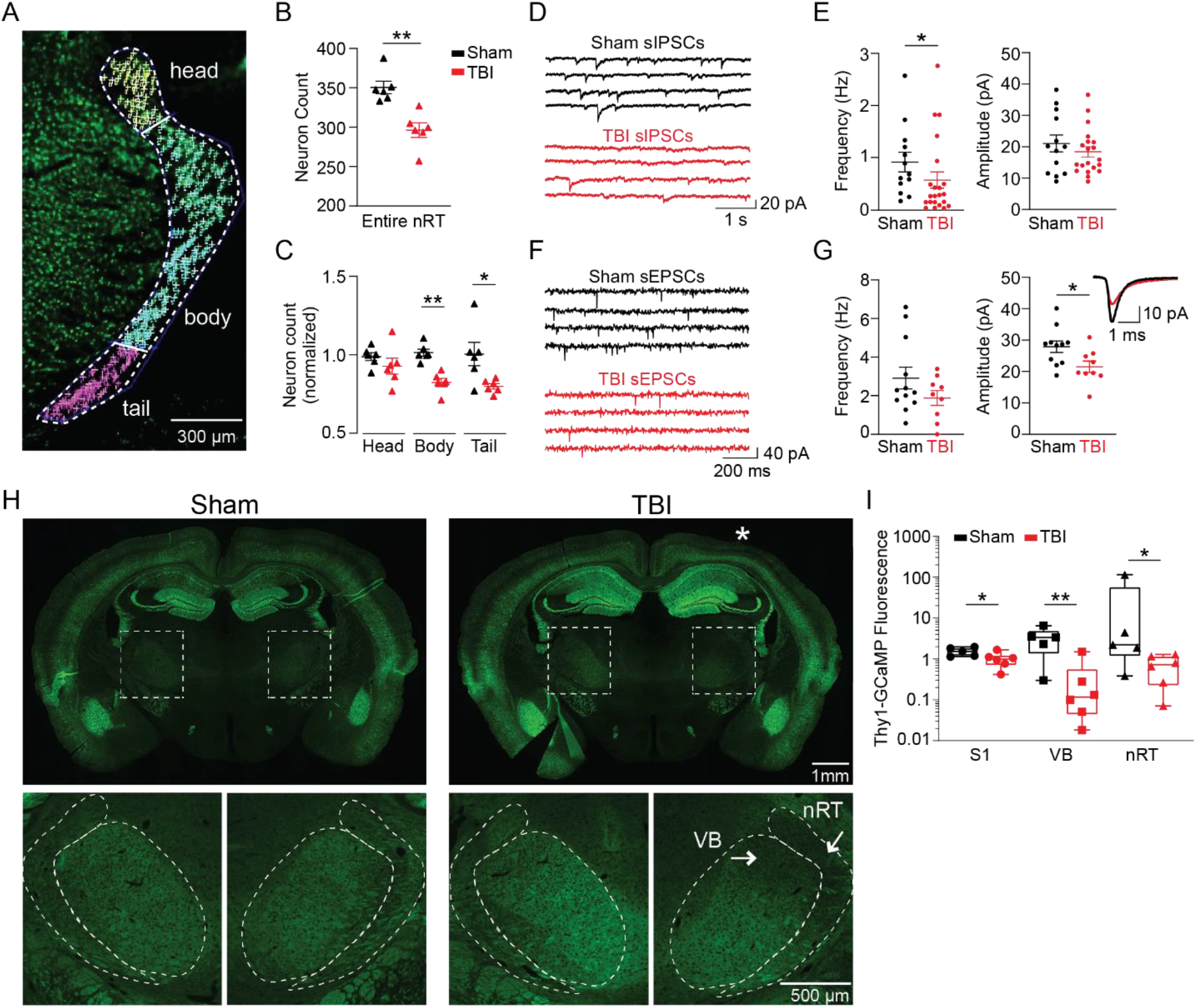
The nRT ipsilateral to the injured cortex shows neuron loss and altered IPSC and EPSC properties three weeks after TBI. A-C) High-magnification coronal image of the nRT showing divisions into “head”, “body”, and “tail” (*50*), and quantification of neuron counts across the entire ipsilateral nRT (B) or per subdivision, normalized to the median value from the sham group (C). Neuron count data represent mean ± SEM, with a Mann-Whitney test and α = 0.05 (*p < 0.05, **p < 0.01). Analysis includes six mice per group (n = three sections per mouse, averaged). D, E) Spontaneous IPSC recordings (D) from representative nRT neurons in sham and TBI mice, and frequency and amplitude distributions (E) in 13 posterior nRT neurons from four sham mice and 22 posterior nRT neurons from six TBI mice. IPSC data represent mean ± SEM analyzed with a Mann-Whitney test and α = 0.05 (*p < 0.05). F, G) Spontaneous EPSC recordings (F) from representative nRT neurons in sham and TBI mice, and frequency and amplitude distributions (G) in 11 posterior nRT neurons from six sham mice and nine posterior nRT neurons from seven TBI mice. Inset shows averaged EPSC traces from single nRT neurons from sham and TBI mice, plotted on the same scale. EPSC data represent mean ± SEM analyzed with a Mann-Whitney test and α = 0.05 (*p < 0.05). H) Representative images of coronal brain sections from Thy1-GCaMP6f mice with sham surgery (left) and TBI (right) (injury site marked by asterisk). Bottom panels show projection terminals from the cortex to VB and nRT. Scale bars, 1 mm (top) and 500 μm (bottom). Reduction in projection terminals from the cortex to VB and nRT (marked by arrows) were observed in n = six TBI mice. I) Quantification of Thy1-GCaMP fluorescence ratios between ipsilateral and contralateral regions in sham and TBI mice. Data represent all points from min to max, with a Mann-Whitney test and α = 0.05 (*p < 0.05, **p < 0.01). Analysis includes five sham mice and six TBI mice (n = three sections per mouse, one image per region).

To test whether C1q might mark functional damage in this circuit, we performed whole-cell patch-clamp recordings in the cortex and thalamus in brain slices at chronic stages of TBI (three to six weeks). We recorded layer-5 pyramidal neurons and fast-spiking GABAergic interneurons in the peri-TBI S1 cortex, glutamatergic neurons in the VB, and GABAergic neurons in the nRT. The neurons’ intrinsic membrane electrical properties and the spontaneous excitatory and inhibitory postsynaptic current (sEPSC and sIPSC) properties were similar between sham and TBI mice in both the peri-TBI cortex and the VB thalamus (see Table S1 for details). However, in the nRT, TBI led to a reduction in the frequency of sIPSCs (Figure 2D-E). Furthermore, nRT sEPSCs were smaller in amplitude, and trended toward a lower frequency (Figure 2F-G). Immunofluorescence staining for GFP in Thy1-GCaMP6f mice, a marker of neuronal calcium levels in corticothalamic neurons, revealed reduced corticothalamic fluorescence in the thalamus after TBI (Figure 2H-I), suggesting that this circuit is indeed impaired.

We conclude that the major long-term effect of TBI on corticothalamic circuits involves disruption of synaptic transmission in the nRT, which coincides with increased C1q expression, reduced cortical inputs, and local neuronal loss. In contrast, neurons in the peri-TBI cortex and the VB appear normal at chronic stages post-TBI (Table S1), suggesting that inflammation - in particular, increased C1q expression - in these regions is not associated with long-term dysfunction in neuronal excitability or synaptic function.

### Blocking C1q function reduces chronic glial inflammation and neuron loss

Increased C1q expression persisted four months post-TBI (Figure S2A-S2B). To test C1q’s causal involvement in the inflammation and neuronal loss observed three weeks post-TBI, we used an antibody that specifically binds to C1q and blocks its downstream activity (*30*). Mice were given i.p. injections of the C1q antibody or a mouse IgG1 isotype control 24 hours after TBI or sham surgery, followed by twice weekly treatments for three weeks (see methods for more details).

TBI mice treated with the anti-C1q antibody showed a strong reduction in inflammation and reduced neuronal loss (Figure 3A-C) relative to control-treated TBI mice, and on average had the same number of nRT neurons as antibody-treated sham mice (Figure 3C). TBI mice treated with the control still showed inflammation and neuron loss three weeks after TBI (Figure 3). As an alternative approach to the antibody treatment, we repeated the study using C1q −/− mice and found that TBI C1q −/− mice also exhibited reduced chronic inflammation and reduced neuron loss in the nRT (Figure S3).

**Figure 3.**
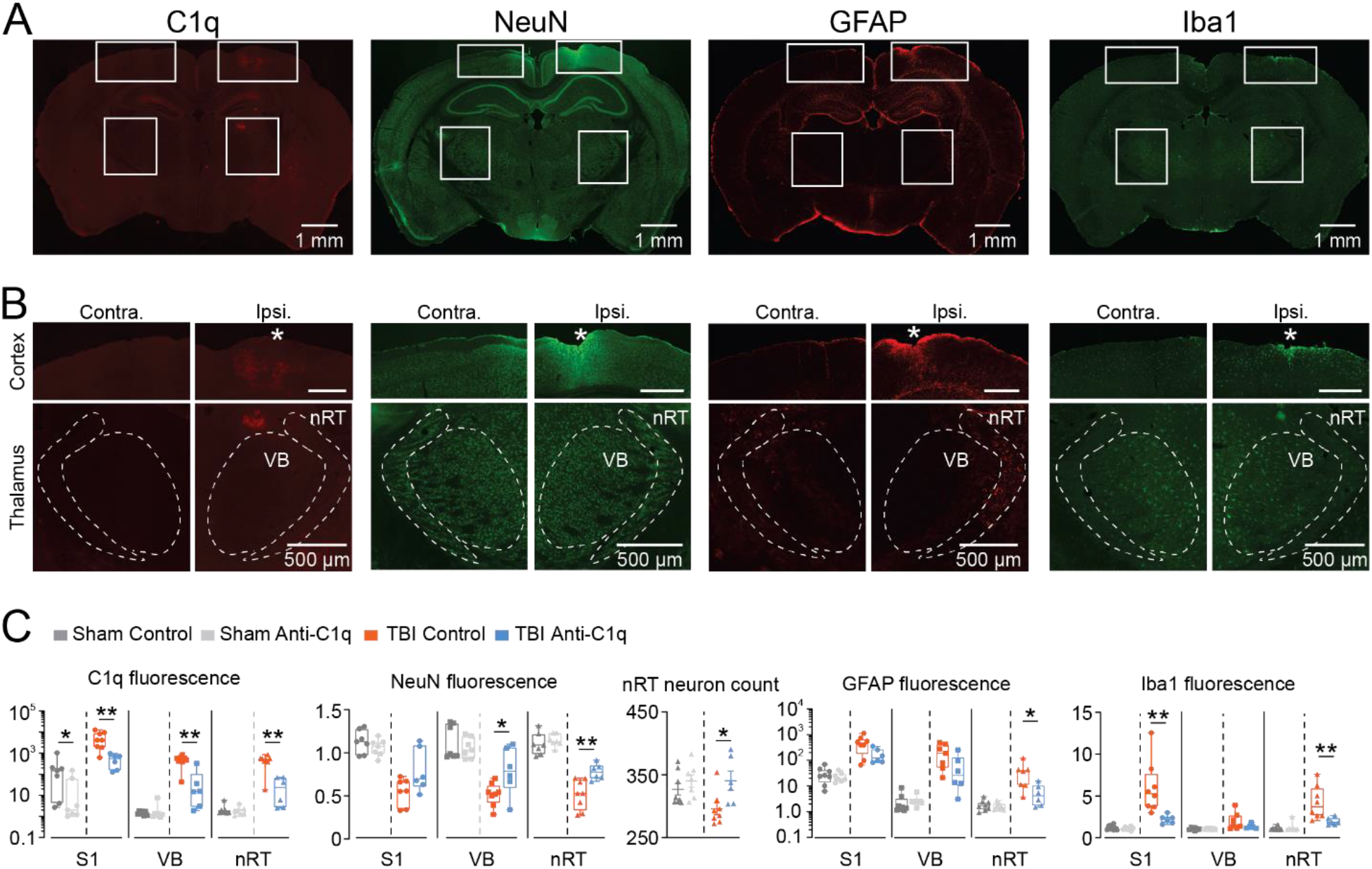
Anti-C1q antibody reduces chronic inflammation and neuron loss three weeks after TBI. A, B) Representative coronal brain sections (A) and close-ups (B) of S1 (top), VB and nRT (bottom) from TBI mice treated with anti-C1q antibody and stained for C1q, NeuN, GFAP, and Iba1. Injury site in the right S1 cortex is marked by an asterisk. Scale bars, 1 mm (A), 500 μm (B). C) Quantification of nRT neuron counts and fluorescence ratios between ipsilateral and contralateral regions in control and antibody-treated sham and TBI mice. Data represent all points from min to max, with a Mann-Whitney test and α = 0.05 (*p < 0.05, **p < 0.01). Analysis includes between six and eight mice per group (n = three sections per mouse, one image per region).

To confirm presence and effects of the anti-C1q antibody in the brain, we measured free drug, free and total C1q, C1s, and albumin levels in naïve, sham and TBI brains after two doses of control or antibody treatment (Figure S4). In plasma from the treated mice, free anti-C1q antibody was observed in both sham and TBI mice treated with the drug (Figure S4A). In agreement, we found that total C1q protein was undetectable in drug-treated animals using an assay that is not affected by free drug. These results suggest that drug-bound C1q is fully cleared from the circulation. Free anti-C1q antibody was observed in treated sham and TBI mice: 0.4-8.6 ug/ml in the ipsilateral side and 0.09-3.8 ug/ml in the contralateral side. The sham and TBI injuries led to a significant increase in ipsilateral C1q and small increase in contralateral C1q in untreated mice. In the anti-C1q treated sham and TBI mice, total C1q levels were significantly reduced in the ipsilateral side and showed trends of reduction in the contralateral side. Measurable levels of free anti-C1q antibody were observed, suggesting that C1q was fully saturated, but not fully cleared as in the periphery.

These outcomes indicate that C1q may lead to inflammation and neuron loss in TBI, and that blocking C1q reduces these deleterious effects.

### TBI leads to long-term changes in cortical states and excitability in freely behaving mice

We next investigated the longitudinal impact of mild TBI, using brain rhythms as a readout of corticothalamic circuit function *in vivo.* To this end, we implanted chronic wireless electrocorticographic (ECoG) devices into sham and TBI mice during the craniotomy/TBI induction surgery, returned mice to their home cages for chronic recording, and analyzed changes in ECoG power over time (Figure 4A-H). We observed a chronic increase in broadband power in TBI during both light epochs (Figure 4C-H) and dark epochs (data not shown).

**Figure 4.**
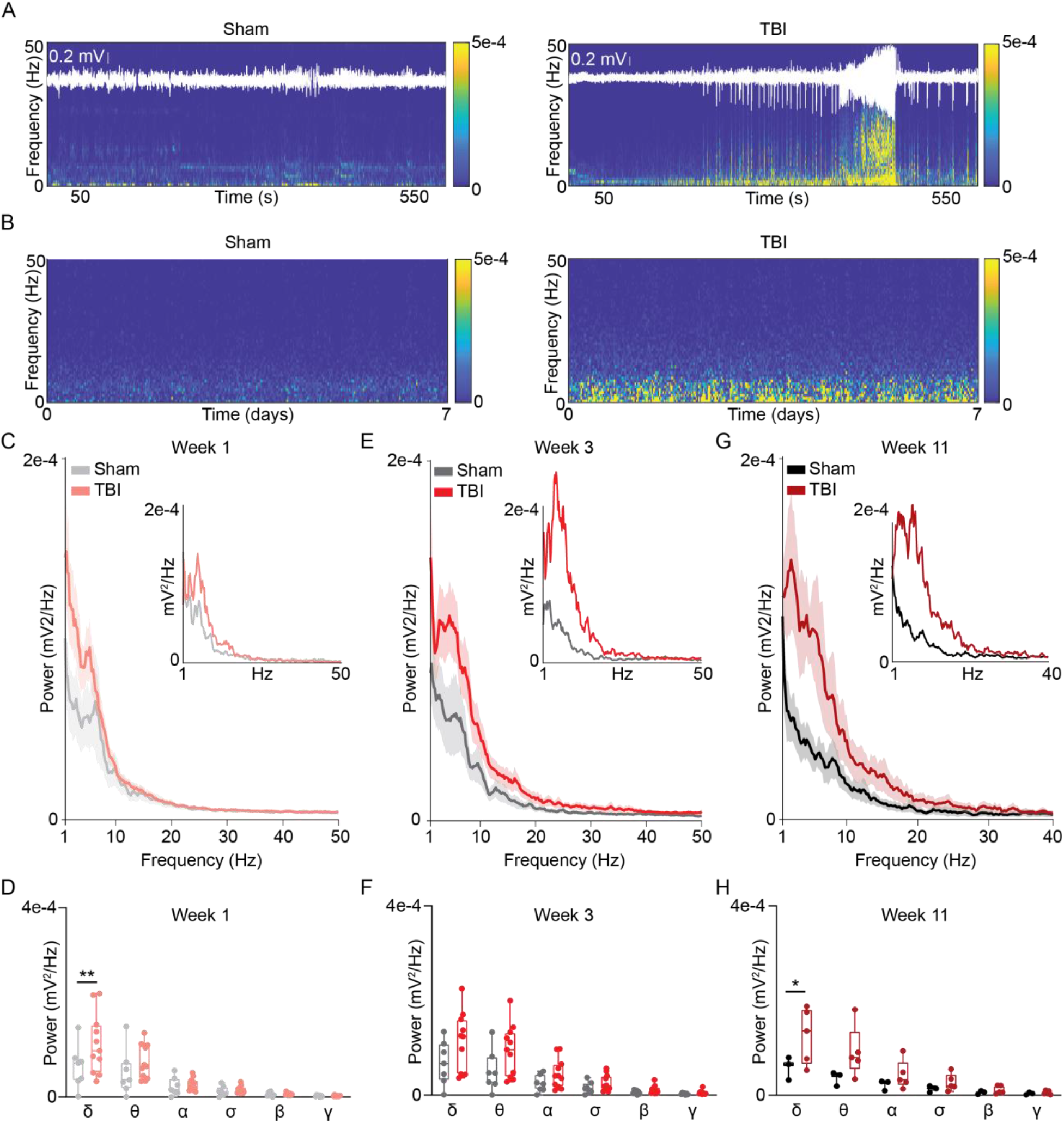
Chronically recorded TBI mice show altered power across different ECoG frequency bands. A) Example 10-minute spectrograms from a sham mouse (left) and TBI mouse (right) taken from the same time point within the first 24 hours of TBI, overlaid with ECoG traces from ipsilateral S1. The TBI spectrogram shows an example of an electrographic seizure, while the sham spectrogram shows normal ECoG activity. Color bar represents power (mV^2^/Hz). B) Example seven-day spectrograms from a sham mouse (left) and TBI mouse (right) showing power across different frequency bands two to three weeks post-TBI. Power bands are sampled every 30 minutes. Color bar represents power (mV^2^/Hz). C) Power spectral density of ECoG activity from sham and TBI cohorts averaged across the first week post-TBI. Inset shows example power spectral density plots from a representative sham and TBI mouse. See methods for details. D) Two-way ANOVAs of average power across frequency bands for the first week post-TBI. Each dot represents power for one mouse. E) Same as C) but at three weeks post-TBI. F) Same as D) but at three weeks post-TBI. G) Same as C) but at 11 weeks post-TBI. H) Same as D) but at 11 weeks post-TBI. Data represent all mice recorded, analyzed with a two-way ANOVA (*p < 0.05, **p < 0.01), even if they died or if the battery ran out before the experimental endpoint. n = eight sham mice, 16 TBI mice. One mouse died within two days post-TBI. The remaining mice were recorded for the first week post-TBI, then recorded for alternating weeks until eleven weeks post-TBI. Delta = 1-4 Hz, theta = 5-8 Hz, alpha = 9-12 Hz, sigma = 13-15 Hz, beta = 16-30 Hz, gamma = 31-50 Hz.

Given that severe TBI has been shown to lead to epileptogenesis over time (*31–32*), we investigated whether mild TBI also resulted in epileptogenesis. We quantified different types of epileptic activities including epileptiform spikes, epileptic discharges, spike-and-wave discharges, and spontaneous focal or generalized seizures at 24 hours and three weeks post-TBI using previously reported classification (*31–32*). In the first 24 hours, 3 out of 16 TBI mice, but none of the 8 sham mice, showed generalized tonic-clonic seizures (GTCSs, Table S2). None of the mice showed GTCSs at later time points (up to three weeks) (Table S2). However, at three weeks post-TBI, we saw more epileptiform spikes in TBI mice (n=9) than in sham mice (n=5), suggesting an increase in excitability (Table S2). Similarly, in another recording setup using simultaneous ECoG and multi-unit thalamic recordings, we observed that TBI mice have spontaneous epileptiform events that include synchronized thalamic bursting and increased normalized theta power, as early as one week and up to three weeks post-TBI (Figure S5).

### Anti-C1q antibody may have modest effects on chronic cortical states in mice with TBI

To determine whether blocking C1q could rescue changes in cortical states, we treated mice with the anti-C1q antibody or isotype control for five weeks, starting 24 hours post-TBI, while maintaining ECoG recordings for up to 9-15 weeks post-TBI (Figure 5A, Figure S6). While the ECoG spectral features were similar within the first week of anti-C1q antibody or control treatment (Figure 5B-C, Figure S6B), analysis of the combined cohorts show that the anti-C1q group trended toward reduced power across most frequency bands at three weeks (Figure 5D-E, Figure S6C).

**Figure 5.**
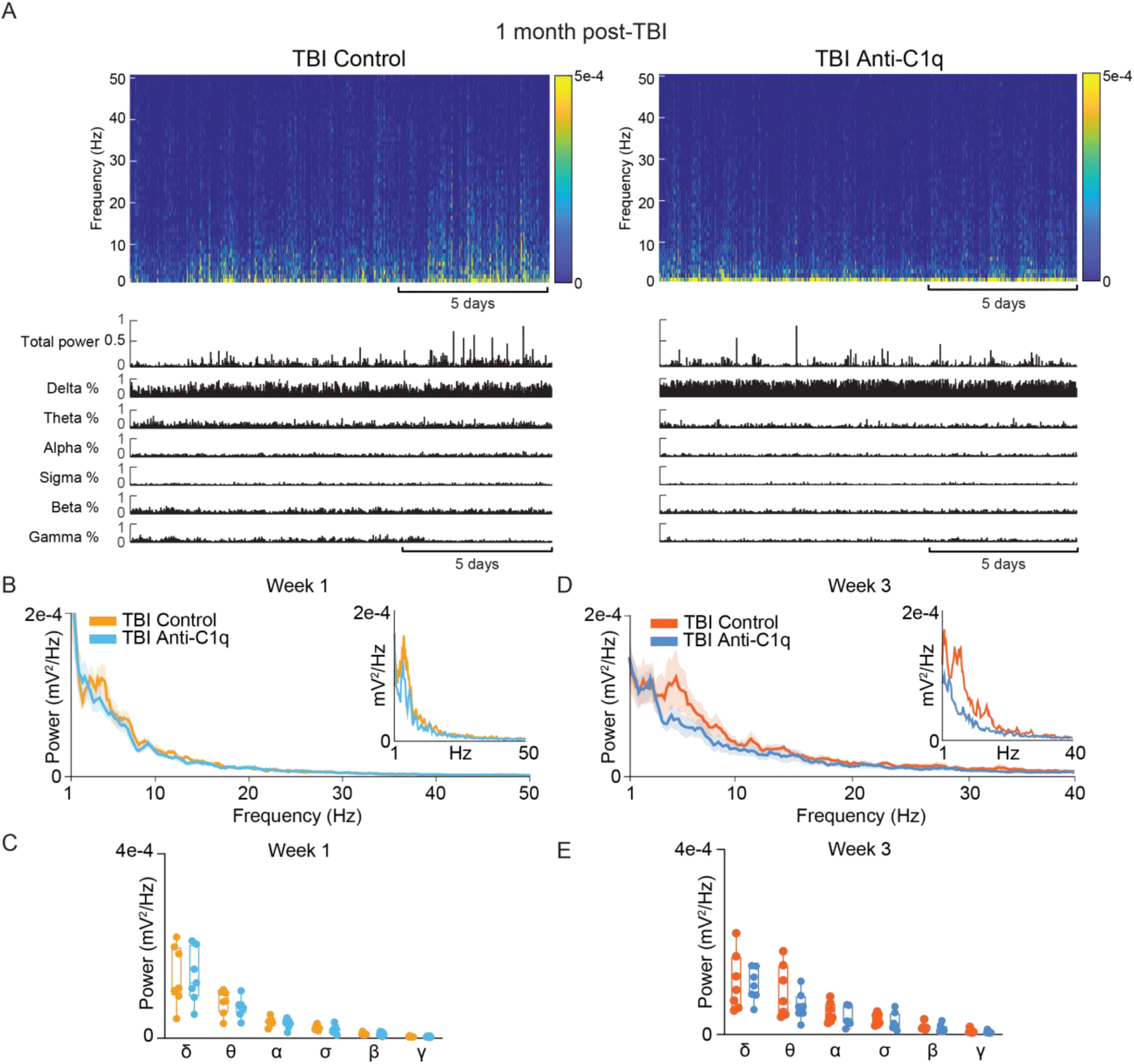
Anti-C1q antibody has modest effects on ECoG spectral features in mice with TBI. A) Example spectrograms (top) and histograms (bottom) from a control-treated mouse (left) and antibody-treated mouse (right) showing power across different frequency bands one month post-TBI. Power bands are sampled every 30 minutes. Color bar represents power (mV^2^/Hz). B) Power spectral density of ECoG activity from control-treated and antibody-treated TBI cohorts averaged across the first week post-TBI. Inset shows example power spectral density plots from a representative control-treated TBI mouse and an antibody-treated TBI mouse. See methods for details. C) Two-way ANOVAs of average power across frequency bands for the first week post-TBI. Each dot represents power for one mouse. D) Same as B) but at three weeks post-TBI. E) Same as C) but at three weeks post-TBI. Data represent all mice recorded, analyzed with a two-way ANOVA, even if they died before treatment ended. n = seven control-treated mice, seven antibody-treated mice. Delta = 1-4 Hz, theta = 5-8 Hz, alpha = 9-12 Hz, sigma = 13-15 Hz, beta = 16-30 Hz, gamma = 31-50 Hz.

Notably, epileptiform activities were not affected by the anti-C1q antibody (Table S2). Three weeks post-TBI, we saw no GTCSs and no differences in the frequency of epileptic events between control-treated and antibody-treated TBI mice (Table S2).

## DISCUSSION

In this study, we set out to understand the role of the C1q pathway in post-TBI secondary injury to the corticothalamic circuit in a mechanistically tractable and highly reproducible mouse model of cortical injury. This model allows us to identify factors such as therapeutic windows, inflammatory phenotypes, and degree of secondary damage, which have been postulated to be important for designing targeted approaches in the treatment of post-TBI outcomes (*13*).

Our study pioneers the use of electrophysiological approaches to study the entire somatosensory corticothalamic circuit after TBI. One powerful tool we employ is chronic ECoG recordings to study the progression of post-traumatic epileptogenesis and changes in cortical rhythms up to four months post TBI. Using electrophysiological approaches at the cellular and circuit levels, we show that TBI alters the synaptic properties of nRT neurons and is associated with increased C1q accumulation that might mediate pathological states in the corticothalamic circuit, including increased broadband activity.

### The nRT as a locus of long-term, secondary impairments post-TBI

We found two kinds of defects in the nRT: neuron loss, and alterations in synaptic properties. nRT neurons degenerated by three weeks after TBI, in agreement with previous observations from the human nRT (*33*), suggesting that even mild cortical injury can lead to neuronal loss in nRT. Potential causes for this neurodegeneration could be loss of cortical inputs causing excitotoxicity in nRT, which has been suggested to be a vulnerable brain region due to high density of axonal afferents from the cortex (*33*). We hypothesized that inflammation plays a major role in this process and that inhibiting inflammation would rescue these defects.

The loss of neurons in the nRT could explain some of the synaptic changes in this area. In particular, we found that three weeks post-TBI, the frequency of IPSCs was reduced in nRT neurons. In many microcircuits, reduced inhibition on GABAergic neurons results in a net increase in inhibition. By contrast, loss of GABAergic inhibition in the nRT results in corticothalamic circuit hyperexcitability, and can even elicit epileptiform activity (*34–35*). Indeed, intra-nRT GABAergic connections are important for coordinating inhibitory output to the excitatory thalamic nuclei and controlling oscillatory thalamic activity (*36*), and their loss is deleterious to the corticothalamic circuit. We speculate that the death of GABAergic neurons in the nRT may contribute to reduced intra-nRT inhibition. This reduced inhibition could cause a loss of feed-forward GABAergic inhibition, which may contribute to increased seizure susceptibility, and increased likelihood of developing post-traumatic epileptic activities.

We also observed deficits in nRT EPSCs, in particular lower amplitude. This alteration is similar to the findings from a mouse model of epilepsy that lacks GluA4 AMPA receptors at the cortico-nRT glutamatergic synapse. This defect results in loss of feed-forward inhibition in the thalamus, and epileptic activities (*37*). We therefore propose that alterations to the nRT EPSCs also contribute to corticothalamic circuit hypersynchrony and seizures, but likely results from a loss of cortical glutamatergic inputs to the nRT after TBI, although loss of other afferents cannot be excluded.

Given that the changes we found in the corticothalamic circuit, and the nRT in particular, have been implicated in epileptic activities and cognitive deficits, our study pinpoints this circuit as a novel potential target for treating long-term TBI outcomes.

Unlike nRT neurons, cortical neurons, such as layer-5 pyramidal neurons and GABAergic fast-spiking interneurons, were not altered by mild TBI at chronic time points. These observations suggest the presence of homeostatic mechanisms that restore or reduce chronic hyperexcitability after TBI in the cortex. They also confirm that at least certain long-term outcomes of TBI must result from nRT dysfunction rather than simply from damage to the cortex. In this regard, it is interesting to see that while cortical neurons appear to have normal excitability and synaptic function at the chronic phase, the cortex shows increased broadband activity, particularly delta. This observation is in agreement with previous magnetoencephalography studies in humans with mild TBI, EEG studies in humans with severe TBI, and EEG studies from rats with severe TBI, which observed increased delta activity at early time points post-TBI (*38–41*). In normal conditions, delta activity is associated with slow wave sleep, quiet wakefulness, and higher cognitive function (*42–43*). In cases of injury, delta waves are associated with a white matter lesion (*39*).

Overall, our findings suggest that the major long-term impact of mild TBI is in the thalamic end of the cortico-thalamo-cortical loop.

### C1q: good or bad? A question of timing

We chose to test the importance of one specific inflammatory pathway, the classical complement pathway, using a pharmacological tool to block C1q in TBI mice. C1q has a well-documented role in normal brain function such as synaptic pruning during development (*24*), as well as its involvement in several neurological diseases (*16–22*). In addition, we had observed that C1q expression was highly increased in the corticothalamic circuit for up to four months after TBI (Figure S2).

Although our mild TBI mice did not develop chronic GTCSs to determine if blocking C1q had an anti-seizure effect, we did observe many other protective effects of the anti-C1q antibody, including reduced inflammation and neurodegeneration. Based on these observations and previous literature implicating differences between protective and harmful inflammatory cell types (*44–45*), we speculate that C1q plays both good and bad roles but at different stages of pathology. At the time of the injury, C1q plays a “beneficial” role, perhaps by aiding with the formation of the glial scar that limits the size of the injury within the primary site of the cortex (*46–47*). However, at the chronic phase, C1q increase plays a maladaptive role in promoting chronic inflammation and secondary neurodegeneration in the nRT.

The cortex also exhibits an increase in C1q, but it does not appear to have a damaging role at this site, or may play a counterbalancing initial protective role since, unlike in the thalamus, the neuronal physiology is similar in the cortex of sham and TBI mice at chronic time points. Our findings suggest the existence of a time window during which the anti-C1q treatment might prevent secondary damage to the thalamus without impairing homeostatic recovery at the cortex.

In conclusion, our study pinpoints C1q as a potential disease modifier that could be targeted for treating devastating outcomes of TBI within a certain time window (in this study, beginning treatment 24 hours post-injury). C1q might also serve as a biomarker to help identify those individuals likely to develop long-term, secondary injuries. Our study also motivates further investigation of the molecular mechanism by which C1q causes neuronal death in nRT, beyond its well-known role in synaptic pruning in health and disease. In addition, by showing that the thalamus is chronically affected by TBI, we identify a potential cause for many TBI-related disabilities such as altered sensory processing, sleep disruption, and epilepsy, and a novel target for post-TBI treatments.

## MATERIALS AND METHODS

### Animals

We performed all experiments per protocols approved by the Institutional Animal Care and Use Committee at the University of California, San Francisco and Gladstone Institutes. Precautions were taken to minimize stress and the number of animals used in each set of experiments. Mice were separately housed after surgical implants. Adult (P30-P180) male CD1 mice were used for most experiments. Adult male Thy1-GCaMP6f mice (Tg(Thy1-GCaMP6f)GP5.17Dkim ISMR_JAX: 025393; C57BL/6 congenic) and C1q null mice (C1qa^tm1Mjw^, ISMR_APB: 1494; C57BL/6 congenic) were used for specific experiments.

### Controlled cortical impact

We anesthetized mice with 2-5% isoflurane and placed them in a stereotaxic frame. We performed a 3 mm craniotomy over the right somatosensory cortex (S1) centered at −1 mm posterior from Bregma, +3 mm lateral from the midline. TBI was performed with a CCI device (Impact One Stereotaxic Impactor for CCI, Leica Microsystems) equipped with a metal piston using the following parameters: 3 mm tip diameter, 15° angle, depth 0.8 mm from the dura, velocity 3 m/s, and dwell time 100 ms. Sham animals received identical anesthesia and craniotomy, but the injury was not delivered.

### Immunostaining and microscopy

We anesthetized mice with a lethal dose of Fatal-Plus and perfused with 4% paraformaldehyde in 1X PBS. Serial coronal sections (30 μm thick) were cut on a Leica SM2000R sliding microtome. Sections were incubated with antibodies directed against C1q (1:700, rabbit, Abcam, ab182451, AB_2732849), GFAP (1:1000, chicken, Abcam, ab4674, AB_304558), GFP (1:500, chicken, Aves Labs, AB_10000240), Iba1 (1:500, rabbit, Wako, 019-19741, AB_839504), and NeuN (1:500, mouse, Millipore, MAB377, AB_2298772) overnight at 4°C. After wash, we incubated sections with Alexa Fluor-conjugated secondary antibodies (1:300, Thermo Fisher Scientific, A-11029) for two hours at room temperature. We mounted sections in an antifade medium (Vectashield) and imaged using a Biorevo BZ-9000 Keyence microscope at 10-20x. Confocal imaging was performed using a confocal laser scanning microscope (LSM880, Zeiss) equipped with a Plan Apochromat 10x/0.45 NA air or 63x/1.4 NA oil immersion objective lens. A multi-line Argon laser was used for 488 nm excitation of AlexaFluor488 and a HeNe laser was used for 561 nm excitation of AlexaFluor594.

### Immunostaining of human tissue

Formalin-fixed, paraffin-embedded tissue was sectioned at 6 μm and mounted on organosilane-coated slides (SIGMA, St. Louis, MO). Representative sections of specimens were processed for hematoxylin/eosin, as well as for immunocytochemistry. Immunocytochemistry for C1q (1:200, rabbit polyclonal; DAKO, Denmark), was carried out on a paraffin-embedded tissue as previously described (*48–49*). Sections were incubated for one hour at room temperature followed by incubation at 4°C overnight with primary antibodies. Single-labeled immunocytochemistry was performed using Powervision method and 3,3-diaminobenzidine as chromogen. Sections were counterstained with hematoxylin. An extensive neuropathological protocol was used (based upon the recommendations of the Brain-Net Europe consortium; Acta Neuropathologica 115(5):497-507 ·2008), including markers such as pTau (AT8), β-Amyloid, pTDP-43 and alpha-synuclein.

### Slice preparation for electrophysiology

We euthanized mice with 4% isoflurane, perfused with ice-cold sucrose cutting solution containing 234 mM sucrose, 11 mM glucose, 10 mM MgSO_4_, 2.5 mM KCl, 1.25 mM NaH_2_PO_4_, 0.5 mM CaCl_2_, and 26 mM NaHCO_3_, equilibrated with 95% O_2_ and 5% CO_2_, pH 7.4, and decapitated. We prepared 250-μm thick horizontal slices for thalamic recordings, and coronal slices for neocortical recordings with a Leica VT1200 microtome (Leica Microsystems). Slices were incubated at 32°C for one hour and then at 24-26°C in artificial cerebrospinal fluid (ACSF) containing 126 mM NaCl, 10 mM glucose, 2.5 mM KCl, 2 mM CaCl_2_, 1.25 mM NaH_2_PO_4_, 1 mM MgSO_4_, and 26 mM NaHCO_3_, and equilibrated with 95% O_2_ and 5% CO_2_, pH 7.4. Thalamic slice preparations were performed as described (*37, 50–51*).

### Patch-clamp electrophysiology

Recordings were performed as previously described (*37, 50–51*). We visually identified S1, nRT, and VB neurons by differential contrast optics with an Olympus microscope and an infrared video camera. Recording electrodes made of borosilicate glass had a resistance of 2.5-4 MΩ when filled with intracellular solution. Access resistance was monitored in all the recordings, and cells were included for analysis only if the access resistance was <25 MΩ. Intrinsic and bursting properties and spontaneous excitatory postsynaptic currents (EPSCs) were recorded in the presence of picrotoxin (50 μM, Sigma) and the internal solution contained 120 mM potassium gluconate, 11 mM EGTA, 11 mM KCl, 10 mM HEPES, 1 mM CaCl_2_, and 1 mM MgCl_2_, pH adjusted to 7.4 with KOH (290 mOsm). We corrected the potentials for −15 mV liquid junction potential.

Spontaneous inhibitory postsynaptic currents (IPSCs) were recorded in the presence of kynurenic acid (2 mM, Sigma), and the internal solution contained 135 mM CsCl, 10 mM EGTA, 10 mM HEPES, 5 mM Qx-314 (lidocaine N-ethyl bromide), and 2 mM MgCl_2_, pH adjusted to 7.3 with CsOH (290 mOsm).

### Surgical implantation of devices for simultaneous recording of ECoG and MUA

The devices for simultaneous ECoG, MUA recordings, and optical manipulations in freely behaving mice were all custom made in the Paz lab as described in (*50–51*). In general, recordings were optimized for assessment of somatosensory subnetworks (S1, somatosensory VB thalamus, and somatosensory nRT).

We implanted cortical screws bilaterally over S1 (contralateral to injury: −0.5 mm posterior from Bregma, −3.25 mm lateral; ipsilateral: +1.0-1.4 mm anterior from Bregma, +2.5-3.0 mm lateral), centrally over PFC (+0.5 mm anterior from Bregma, 0 mm lateral), and in the right hemisphere over V1 (−2.9 mm posterior from Bregma, +2.7 mm lateral). For MUA recordings in VB, we implanted electrodes at −1.65 mm posterior from Bregma, +1.75 mm lateral, with the tips of the optical fiber at 3.0 mm and two electrodes at 3.25 mm and 3.5 mm ventral to the cortical surface. For MUA recordings in nRT, we implanted electrodes at −1.4 mm posterior from Bregma, +2.1 mm lateral, with the tips of the optical fiber at 2.7 mm and two electrodes at 2.9 mm, and 3.0 mm ventral to the cortical surface, respectively.

### In vivo electrophysiology and behavior

Non-chronic MUA electrophysiological recordings in freely behaving mice were performed as described using custom-made optrode devices (*50–51*). ECoG and thalamic LFP/MU signals were recorded using RZ5 (TDT) and sampled at 1221 Hz, with thalamic MUA signals sampled at 24 kHz. A video camera that was synchronized to the signal acquisition was used to continuously monitor the animals. We briefly anesthetized animals with 2% isoflurane at the start of each recording to connect for recording. Each recording trial lasted 15-60 min. To control for circadian rhythms, we housed our animals using a regular light/dark cycle, and performed recordings between roughly 9:00 am and 6:00 pm. All the recordings were performed during wakefulness. We validated the location of the optrodes by histology after euthanasia in mice that did not experience sudden death and whose brains we were able to recover and process.

### Surgical implantation of devices for chronic ECoG recordings

The wireless telemetry devices we used for chronic ECoG recordings were purchased from Data Sciences International (DSI). After performing controlled cortical impact surgery, we implanted cortical screws bilaterally over S1 as described above. The battery/transmitter device was placed under the skin over the right shoulder. We began recording mice as soon as they recovered from the surgery. Mice were singly housed in their home cages, which were placed over receivers that sent signals to an acquisition computer. ECoG signals were continuously recorded from up to eight mice simultaneously using Ponemah software (DSI), and sampled at 500 Hz.

### Statistical analyses

All numerical values are given as means and error bars are standard error of the mean (SEM) unless stated otherwise. Parametric and non-parametric tests were chosen as appropriate and were reported in figure legends. Data analysis was performed with MATLAB (SCR_001622), GraphPad Prism 7/8 (SCR_002798), ImageJ (SCR_003070), Ponemah/NeuroScore (SCR_017107), pClamp (SCR_011323), and Spike2 (SCR_000903).

### Image analysis and cell quantification

We selected regions of interest (ROIs) for S1, nRT, and VB from 10x Keyence microscope images opened in ImageJ (SCR_003070). To ensure that each ROI covered the same area on the ipsilateral and contralateral sides of the injury site, the first ROIs were duplicated and repositioned over the opposite hemisphere. The image was then converted to 8-bit. The upper threshold was adjusted to the maximum value of 255, and the lower threshold was increased from 0 until the pixel appearance most closely matched the fluorescence staining from the original image. A particle analysis was run on the ROIs using the same threshold boundaries for all sections with the same stain. An integrated density ratio was calculated for each brain region by dividing the ipsilateral integrated pixel density by the contralateral integrated pixel density. The integrated density ratios from three sections per animal were averaged to get a single average ratio per brain area for each animal.

nRT cell counts were performed on sections stained with NeuN. The nRT was outlined in ImageJ (SCR_003070) and we performed a manual cell count of neuronal cell bodies using the manual counter plugin.

### Analysis of electrophysiological properties

The input resistance (R_in_) and membrane time constant (τ_m_) were measured from the membrane hyperpolarizations in response to low intensity current steps (−20 to −60 pA). The reported rheobase averages and SEMs were calculated based on the current which first caused at least one action potential during the stimulus per recording. All data from Table S1 were analyzed using a Mann-Whitney test with α = 0.05 (*p < 0.05, **p < 0.01, ***p < 0.001, ****p < 0.0001), using GraphPad Prism 7 (SCR_002798).

Cumulative probability distributions were generated in MATLAB (SCR_001622) from 11 sham nRT neurons and 9 TBI neurons, using 200 randomly selected events from each cell.

### ECoG spectral and seizure analysis

Spectrograms were generated for frequencies between 1 and 50 Hz using the short-time Fourier transform with 0.5 s Hamming windows and 98% overlap between segments. Spectrograms and cumulative distribution functions were generated in MATLAB. ECoG frequency bands were divided as follows: delta = 1-4 Hz, theta = 5-8 Hz, alpha = 9-12 Hz, sigma = 13-15 Hz, beta = 16-30 Hz, gamma = 31-50 Hz. Fast Fourier transforms were generated in MATLAB. Epileptic activities were analyzed manually within the time windows specified for each experiment. Epileptic discharges (EDs) were defined as clusters of interictal spiking events lasting five seconds or longer. Spike-and-wave discharges (SWDs) were defined as symmetrical 6-8 Hz spiking events lasting two seconds or longer. Generalized tonic-clonic seizures (GTCSs) were defined as spiking events lasting at least 30 seconds and present in both ECoG channels. Abnormal events were defined as any irregular spiking events that did not fit into any of the other categories.

### Anti-C1q antibody

The anti-C1q is antibody ANX-M1, and the control is a mouse isotype IgG1 antibody (Annexon Biosciences). All mice were treated at a concentration of 100 mg/kg, and previous studies have reported no toxicity in rodents. The antibody has been characterized as described in (*21, 30, 52*). Mice were first treated 24 hours post-TBI, and continued receiving treatment every three days (four days post-TBI, seven days post-TBI, etc.) for three weeks, or until the experimental end point as noted in Figure 5.

### Treatment paradigm and tissue lysis

For the PK study, mice underwent sham or TBI surgery on Day 0, and were treated intraperitoneally with 100 mg/kg of anti-C1q antibody (M1) or isotype control on Day 1 and 4. Mice were perfused with PBS on Day 5. Plasma and brains (ipsi- and contra-lateral sides) were collected and flash frozen. Brains (without olfactory bulb and cerebellum) were lysed in 1:10 w/v BupH™ Tris Buffered Saline (Thermo Scientific 28379) + protease inhibitor cocktail (Thermo Scientific A32963) by homogenizing with 7 mm steel bead in Qiagen TissueLyser for two minutes at 30 Hz. Lysates were then spun at 17,000 x g for 20 minutes. Supernatants were used for ELISA assays.

### Pharmacokinetic (PK) and pharmacodynamic (PD) ELISA assays

The levels of free anti-C1q drug M1 (PK), free C1q, total C1q, C1s and Albumin were measured using sandwich ELISAs. Black 96 well plates (Nunc 437111) were coated with 75 uL of respective capture protein/antibody: human C1q protein for PK (complement Tech), mouse monoclonal anti-C1q (Abcam, ab71940) for C1q-free, rabbit polyclonal anti-C1q (Dako, A0136) and rabbit polyclonal anti mouse C1s (LSBio, C483829) for C1s, in bicarbonate buffer (pH 9.4) overnight at 4°C. Next day, the plates were washed with dPBS pH 7.4 (Dulbecco’s phosphate-buffered saline) and blocked with dPBS containing 3% bovine serum albumin (BSA). Standard curves were prepared with purified proteins in assay buffer (dPBS containing 0.3% BSA and 0.1% Tween20). Samples were prepared in the assay buffer at appropriate dilutions. The blocking buffer was removed from the plate by tapping. Standards and samples were added at 75 uL per well in duplicates and incubated with shaking at 300 rpm at room temperature for one hour for PK measurements. For complement assays, samples were incubated overnight at 4°C followed by 37°C for 30 minutes and then room temperature for one hour. Plates were then washed three times with dPBS containing 0.05% Tween20 and 75 uL of alkaline-phosphatase conjugated secondary antibodies (goat anti-mouse IgG for PK, M1 for C1q free, rabbit polyclonal anti-C1q for C1q total, rabbit polyclonal anti-C1s for C1s) were added to all wells. Plates were incubated at room temperature with shaking for one hour, washed three times with dPBS containing 0.05% Tween20 and developed using 75 uL of alkaline phosphatase substrate (Life Technologies, T2214). After 20 minutes at room temperature, plates were read using a luminometer. Albumin assay was done using a matched antibody pair from Abcam (ab210890), followed by Avidin-AP secondary antibody for detection. Standards were fit using a 4PL logistic fit and concentration of unknowns determined. Analyte levels were corrected for dilution factors.

#### SUPPLEMENTAL FIGURES

**Figure S1.**
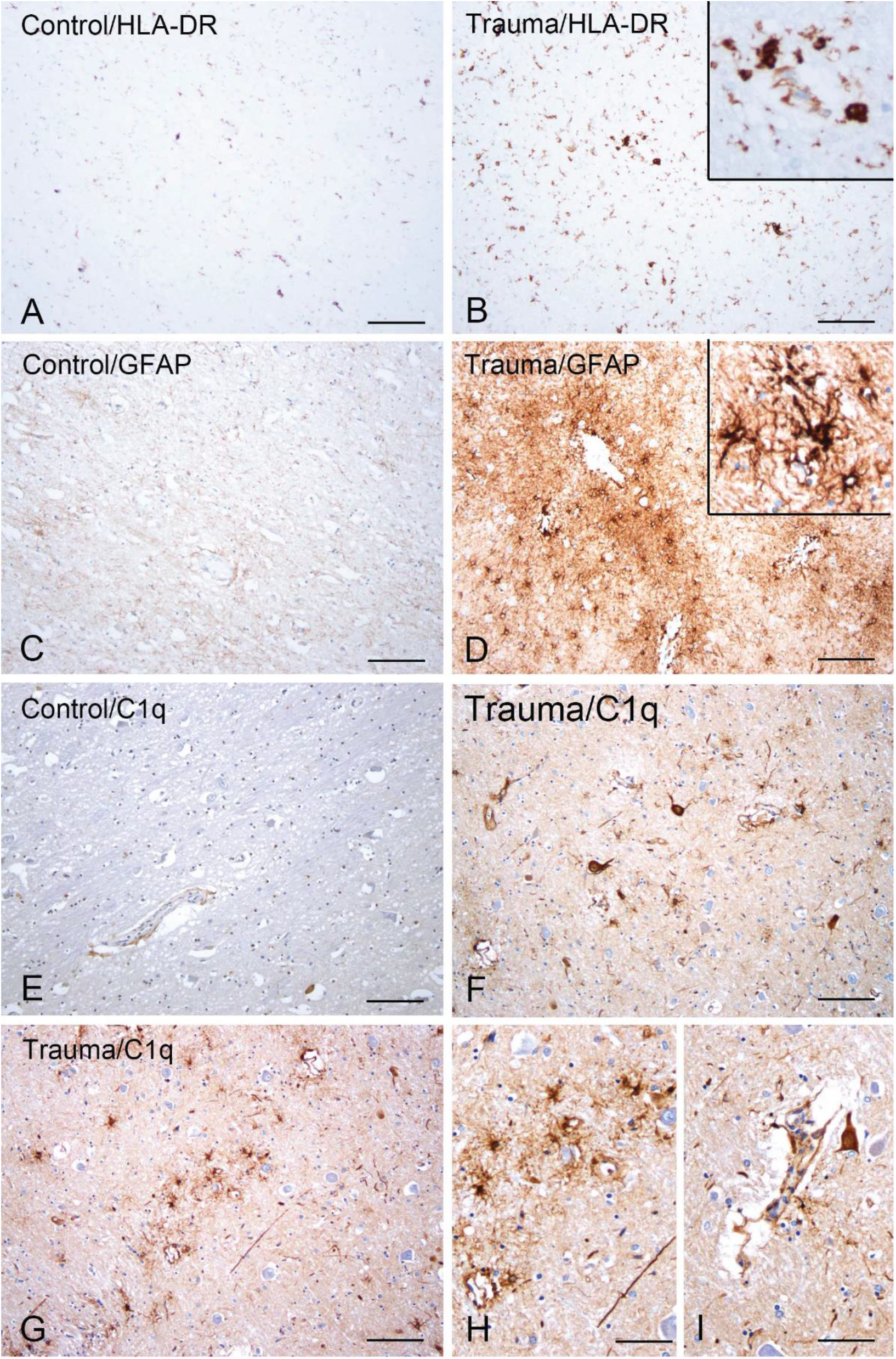
Postmortem brain tissue from a patient with TBI shows chronic inflammation eight days after TBI. A) Postmortem brain tissue from one control patient stained for HLA-DR, a marker for an MHC class II cell surface receptor that is expressed in microglia and macrophages. Case information: male, age 78. Scale bar, 1 cm. B) Postmortem brain tissue from one TBI patient stained for HLA-DR. Case information: male, age 79; fall accident, Injury Severity (GCS): moderate, CT: cerebral edema; no epilepsy (post-TBI: eight days); no history of neurological diseases and without evidence of cognitive decline, based on the last clinical evaluation; no evidence of primary neurodegenerative pathology, evidence of trauma-induced diffuse axonal damage. Scale bar, 1 cm. C) Same as (A) but stained for GFAP. Scale bar, 1 cm. D) Same as (B) but stained for GFAP. Scale bar, 1 cm. E) Same as (A) but stained for C1q. Scale bar, 1 cm. F-I) Same as (B) but stained for C1q. Scale bars, 1 cm (F-G) and 40 μm (H-I).

**Figure S2.**
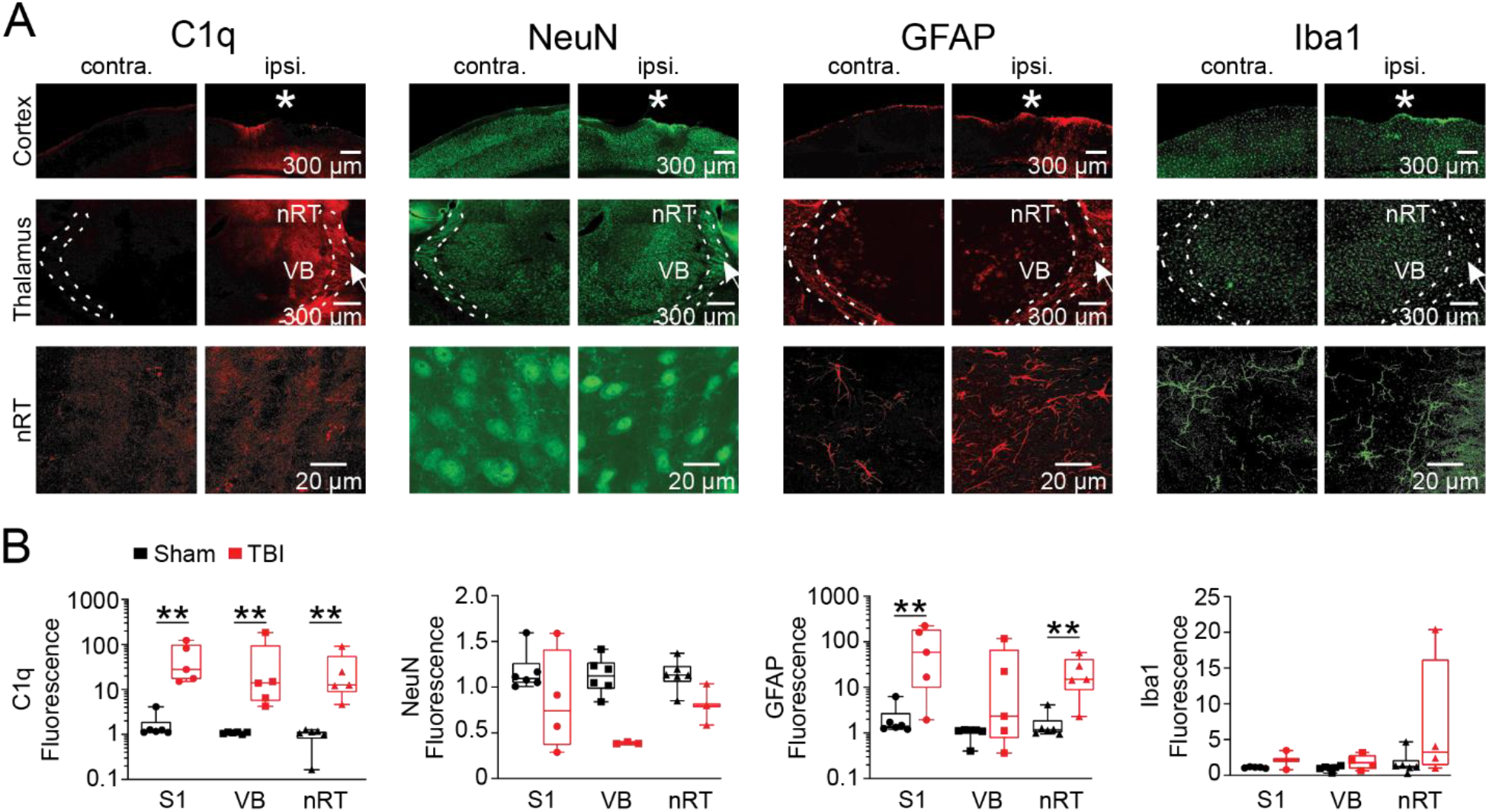
The injured cortex and functionally connected thalamus show chronic inflammation and neuron loss four months after TBI. A) Close-up images of S1 (top), VB and nRT (middle), and confocal images of nRT (bottom), stained for C1q, NeuN, GFAP, and Iba1. Injury site in the right S1 cortex is marked by an asterisk. Arrow in nRT indicates location of confocal image. Scale bars, 300 μm (top/middle) and 20 μm (bottom). B) Quantification of fluorescence ratios between ipsilateral and contralateral regions in sham and TBI mice. Data represent all points from min to max, with a Mann-Whitney test and α = 0.05 (*p < 0.05, **p < 0.01). Analysis includes between four and six mice per group (n = one to three sections per mouse, one image per region).

**Figure S3.**
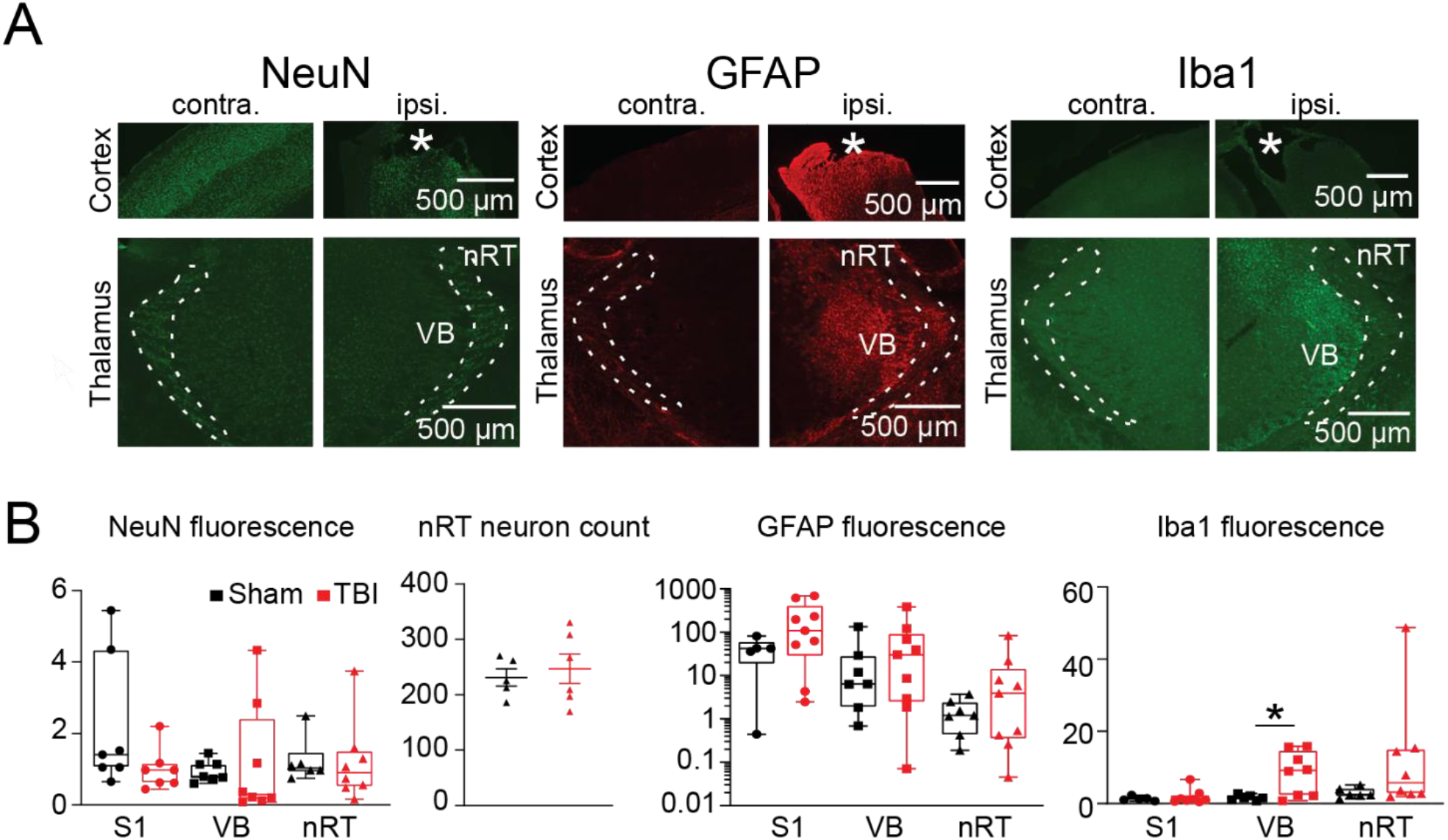
C1q −/− mice show reduced inflammation and neuron loss three weeks after TBI. A) Close-up images of S1 (top), VB and nRT (bottom) stained for NeuN, GFAP, and Iba1. Injury site in the right S1 cortex is marked by an asterisk. Scale bars, 500 μm. B) Quantification of fluorescence ratios between ipsilateral and contralateral regions and nRT neuron counts in sham and TBI C1q −/− mice. Data represent all points from min to max, with a Mann-Whitney test and α = 0.05 (*p < 0.05, **p < 0.01). Analysis includes between four and six mice per group (n = one to three sections per mouse, one image per region).

**Figure S4.**
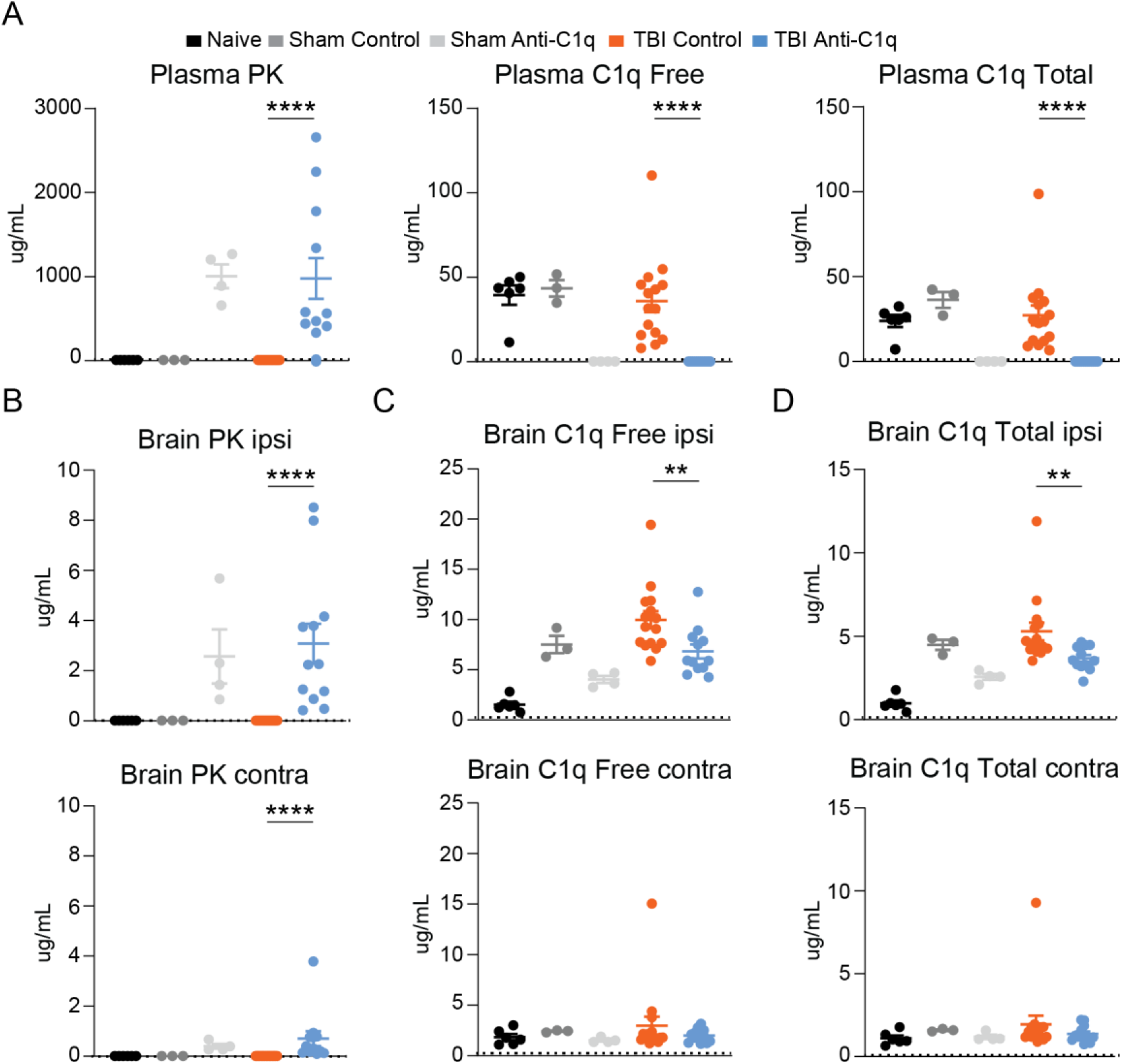
Plasma and brain PK/PD show presence of free drug and reduced C1q in anti-C1q drug-treated sham and TBI mice. A) Plasma levels of free drug, C1q-free, and C1q-total were measured using sandwich ELISAs after TBI and sham mice were treated with two doses of 100 mg/kg anti-C1q or isotype control antibodies. Dotted line shows lower limit of quantification. B-F) Levels of free drug (B), C1q-free (C), and C1q-total (D) were measured in brain lysates in the ipsilateral (top) and contralateral (bottom) sides using sandwich ELISAs. Naïve mice were negative controls. Dotted line shows lower limit of quantification. Data represent all points from min to max, with a Mann-Whitney test between TBI control and TBI anti-C1q, and α = 0.05 (*p < 0.05, **p < 0.01, ***p < 0.001, ****p < 0.0001). Analysis includes between three and 15 mice per group.

**Figure S5.**
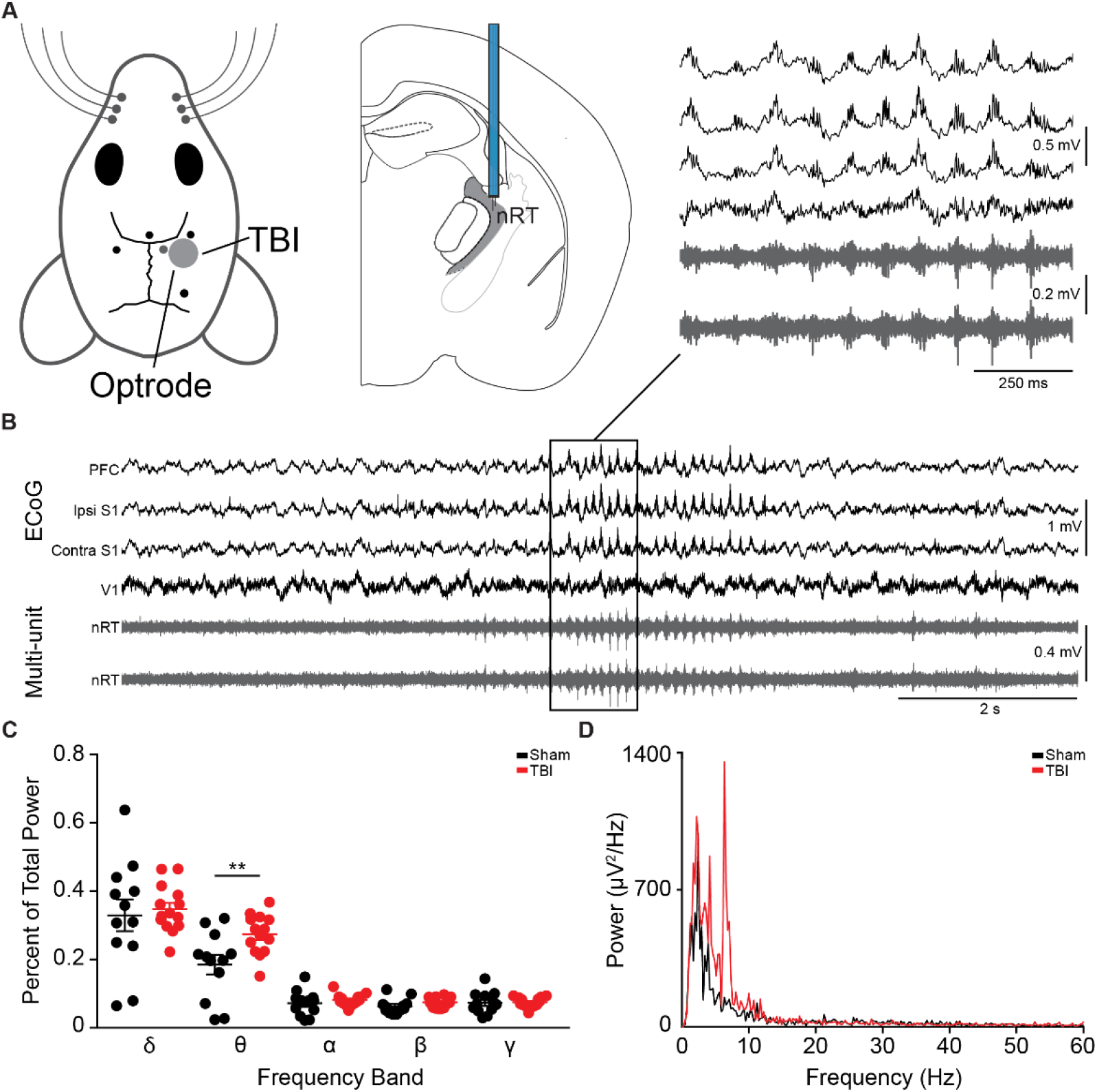
Mice with TBI have spontaneous seizure-like events in the theta to alpha frequency range that are time-locked with thalamic bursting. A) Diagram of recording locations for *in vivo* experiments. Left, ECoG recording sites and TBI location are shown on the mouse skull. Right, approximate location of tungsten depth electrodes implanted unilaterally in the nRT. B) Representative ECoG traces from cortical recording sites and multi-unit traces from nRT showing a spontaneous seizure-like event. C) Power spectral analysis showing the average power across different frequency bands in the first 15 minutes of baseline ECoG signal from the ipsilateral S1 cortex in sham and TBI mice. D) Periodogram showing the power across frequencies taken from the first 15 minutes of baseline ECoG signal from the ipsilateral S1 cortex in a representative sham and TBI mouse. Data represent mean ± SEM analyzed with a Mann-Whitney test and α = 0.05 (*p < 0.05, **p < 0.01). Analysis includes between 12 and 14 mice per group.

**Figure S6.**
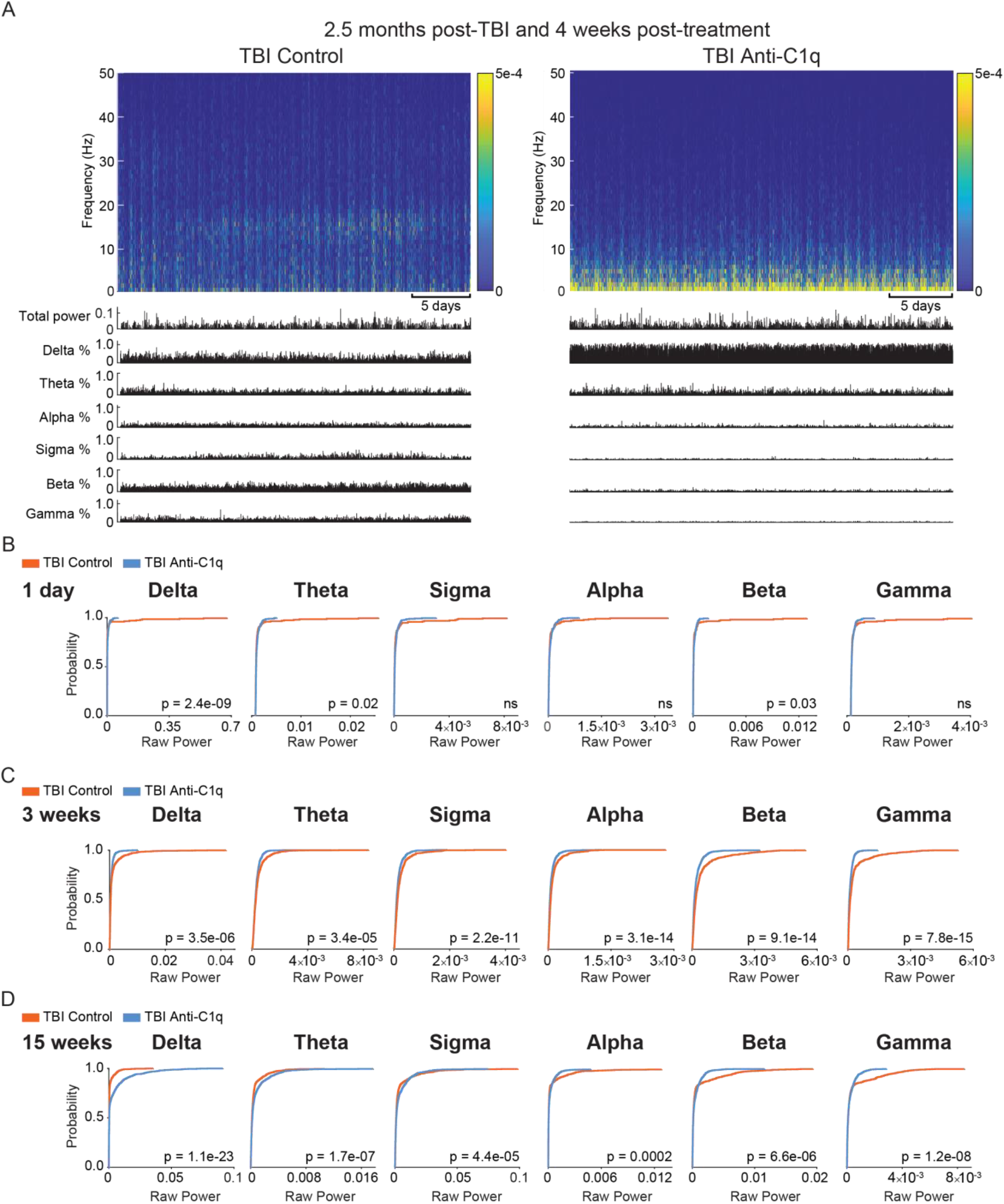
Anti-C1q antibody has chronic disease-modifying effects on ECoG power in mice with TBI. A) Example spectrograms (top) and histograms (bottom) from a control-treated mouse (left) and antibody-treated mouse (right) showing power across different frequency bands 2.5 months post-TBI, which was four weeks after the treatment ended. Power bands are sampled every 30 minutes. B) Cumulative distribution functions for control-treated and antibody-treated cohorts sampled across different frequency bands in the first day post-TBI. We sampled 48 points from the first 24 hours within the start of each recording. C) Same as B, but at three weeks post-TBI. We sampled 232 points between 15.25-20.1 days from the start of each recording. D) Same as B, but at 9-15 weeks post-TBI. We sampled 296 points between 104.6 to 110 days from the start of each recording. Data represent all mice recorded, even if they died before treatment ended. One control-treated mouse and one antibody-treated mouse died within three weeks post-TBI, two control-treated mice died within six weeks post-TBI, and the remaining mice were recorded for at least nine weeks post-TBI. At 24 hours, n = seven control-treated mice, seven antibody-treated mice. At three weeks, n = seven control-treated mice, seven antibody-treated mice. At 9-15 weeks n = six control-treated mice, four antibody-treated mice. Delta = 1-4 Hz, theta = 5-8 Hz, alpha = 9-12 Hz, sigma = 13-15 Hz, beta = 16-30 Hz, gamma = 31-50 Hz. ns = p > 0.05.

**Table S1.**
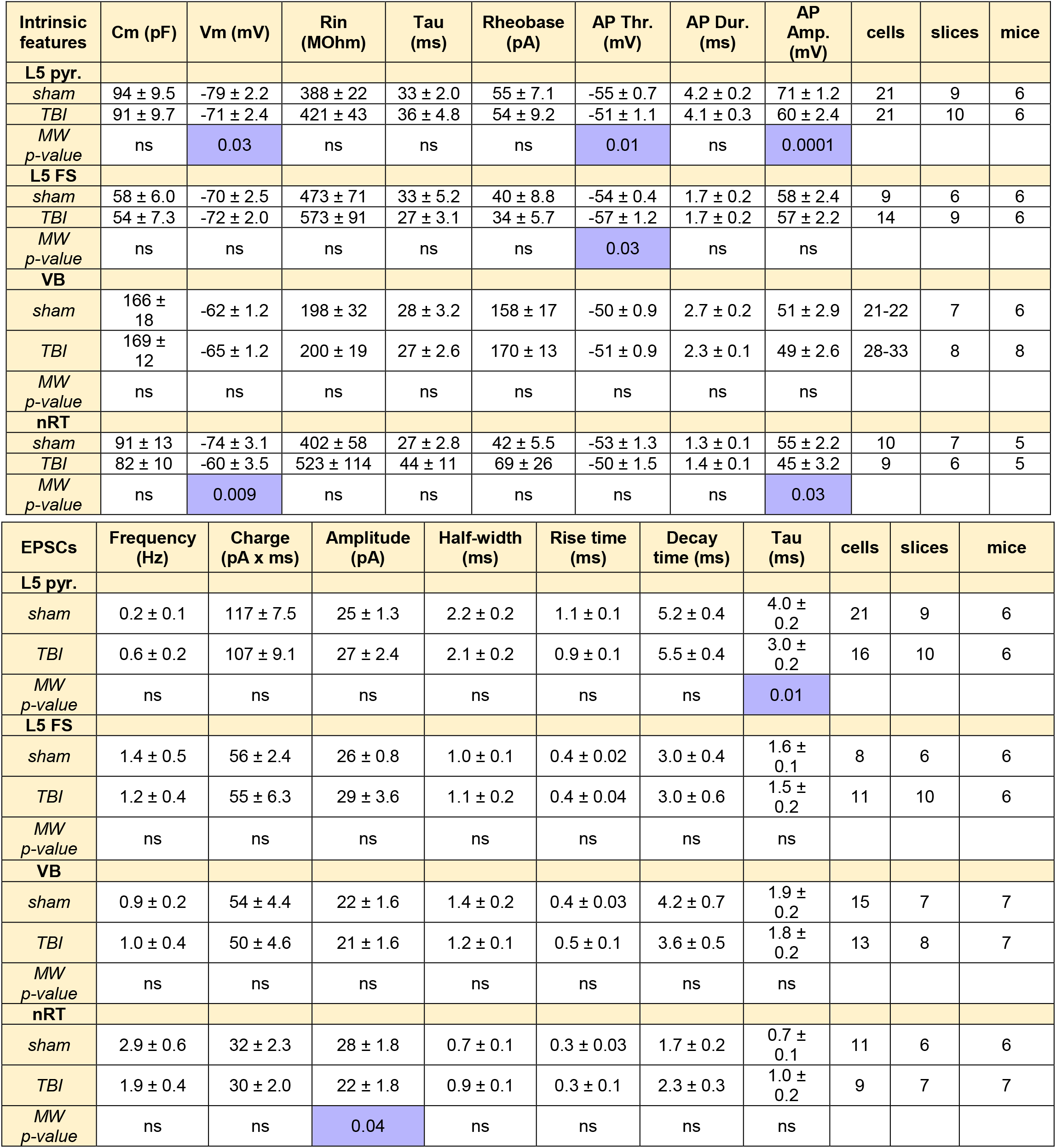
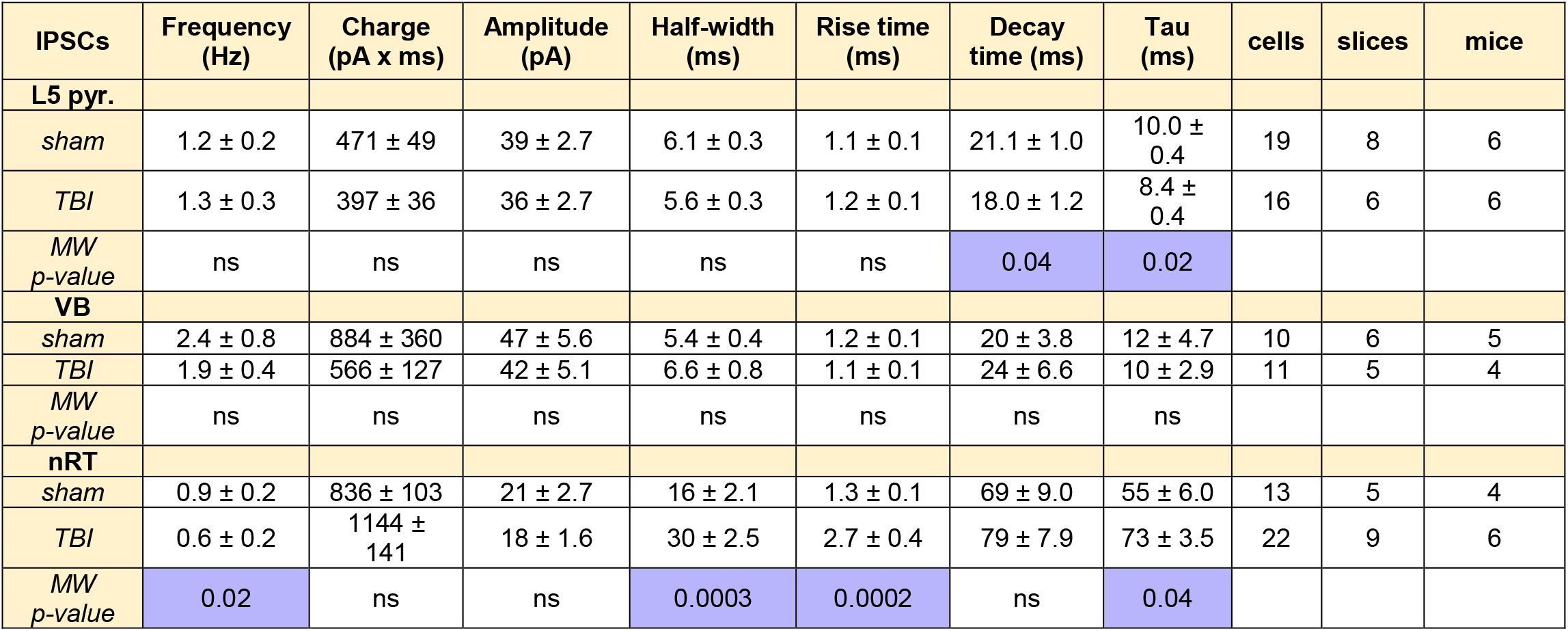
Summary of intrinsic properties, EPSC, and IPSC data recorded from S1 cortex, VB, and nRT. Mice were recorded between three and six weeks post-TBI, and recording conditions are described in the patch-clamp electrophysiology section of the methods. A Mann-Whitney test was performed for statistical analysis.

**Table S2.** Summary of epileptiform activity analysis in sham, TBI, control-treated TBI, and antibody-treated TBI mice. Mice were recorded continuously starting the day of the TBI up until several weeks post-TBI. Surgical and recording conditions are described in the methods section titled “Surgical implantation of devices for chronic ECoG recordings”. Analysis was performed on the first 24 hours post-TBI, and across a 48 hour window at three weeks post-TBI. A repeated measures mixed-effects ANOVA was performed for statistical analysis.

| First 24 hours | Epileptiform spikes | Epileptiform discharges | Spike-and-wave discharges | Generalized tonic-clonic seizures | Acute post-injury mortality |
| --- | --- | --- | --- | --- | --- |
| *Sham* | 8/8 (100%) | 6/8 (75%) | 0/8 (0%) | 0/8 (0%) | 0/8 (0%) |
| *TBI* | 16/16 (100%) | 13/16 (81%) | 4/16 (25%) | 3/16 (19%) | 0/16 (0%) |
| **Drug study** |  |  |  |  |  |
| *TBI Vehicle* | 7/7 (100%) | 5/7 (71%) | 0/7 (0%) | 2/7 (28%) | 0/7 (0%) |
| *TBI anti-C1q* | 7/7 (100%) | 5/7 (71%) | 0/7 (0%) | 1/7 (14%) | 0/7 (0%) |
| 3 weeks | Epileptiform spikes | Epileptiform discharges | Spike-and-wave discharges | Generalized tonic-clonic seizures |  |
| *Sham* | 7/7 (100%) | 1/7 (14%) | 0/7 (0%) | 0/7 (0%) |  |
| *TBI* | 11/11 (100%) | 3/11 (27%) | 1/11 (9%) | 0/11 (0%) |  |
| **Drug study** |  |  |  |  |  |
| *TBI Vehicle* | 7/7 (100%) | 2/7 (28%) | 0/7 (0%) | 0/7 (0%) |  |
| *TBI anti-C1q* | 7/7 (100%) | 3/7 (43%) | 0/7 (0%) | 0/7 (0%) |  |
|  | Epileptiform spikes | Epileptiform discharges | Spike-and-wave discharges | Generalized tonic-clonic seizures | mice |
| *Sham – 24h* | 234 ± 62 | 4 ± 2 | 0 | 0 | 8 |
| *TBI – 24h* | 452 ± 178 | 8 ± 6 | 1 ± 0.6 | 0.4 ± 0.2 | 16 |
| *Sham – 3wk* | 66 ± 38 | 0.7 ± 0.7 | 0 | 0 | 7 |
| *TBI – 3wk* | 292 ± 114 | 2 ± 1 | 0.09 ± 0.09 | 0 | 11 |
| *Mixed-effects analysis* | ns | ns | ns | ns |  |
| **Drug study** |  |  |  |  |  |
| *TBI Vehicle – 24h* | 278 ± 79 | 4 ± 1 | 0 | 0.6 ± 0.4 | 7 |
| *TBI anti-C1q – 24h* | 137 ± 55 | 1 ± 0.5 | 0 | 0.3 ± 0.3 | 7 |
| *TBI Vehicle – 3wk* | 300 ± 92 | 0.3 ± 0.2 | 0 | 0 | 7 |
| *TBI anti-C1q – 3wk* | 274 ± 50 | 1 ± 0.8 | 0 | 0 | 7 |
| *Mixed-effects analysis* | ns | ns | ns | ns |  |

## Acknowledgments

We thank Annexon Biosciences for providing the antibodies. Thanks to Meredith Calvert and the Gladstone Histology & Light Microscopy Core for help with confocal microscopy and ROI analysis. Thanks to Jasper Anink for help with histology on human tissue. Thanks to lab managers Scott Brovarney, Juan Alacauter, and Irene Lew for mouse colony management, and thanks to Françoise Chanut and other editors for critical feedback on the manuscript.

## Author contributions

SSH and JTP conceptualized the project, and wrote and edited the manuscript. SSH administered and/or carried out all experiments including investigation, analysis, and visualization. OA contributed to the anti-C1q histology study and did antibody treatments, histological staining, imaging, and image analysis. BH contributed to setup and data analysis for the chronic ECoG recordings. FSC helped with surgeries and data analysis for the chronic ECoG recordings. AHC and ARM contributed to histological staining and imaging. SS, YAZ, and TY provided the anti-C1q antibodies and gave feedback on all anti-C1q experiments. VM, LJK, and PS carried out all PK/PD ELISA assays. EA provided histological staining on postmortem brain tissue from human patients with TBI. JTP provided supervision and funding.

